# Distinct immunity protein families mediate compartment-specific neutralisation of a bacterial toxin

**DOI:** 10.1101/2025.05.31.657152

**Authors:** Felicity Alcock, Yaping Yang, Justin Deme, Guillermina Casabona, Chriselle Mendonca, Fatima Ulhuq, Susan Lea, Tracy Palmer

## Abstract

*Staphylococcus aureus* utilises a type VII secretion system (T7SS) to secrete antibacterial toxins that target competitor bacteria. EsxX is a T7SS substrate protein secreted by ST398 strains that has an LXG domain at its N-terminus. Here we show that the EsxX C-terminus is a membrane-depolarising toxin with a glycine zipper motif. EsxX is profoundly toxic to bacteria, displaying toxicity from both cytoplasmic and extracellular compartments. A pair of polytopic membrane proteins, ExiCD, protect cells from intoxication by extracellular EsxX. By contrast, a distinct soluble heterodimer, ExiAB, neutralises cytoplasmic EsxX by sequestration of its glycine zipper motif in a binding groove on ExiB. The *exiA-exiB* gene pair co-occur in staphylococcal genomes with *esxX*, invoking a model whereby ExiAB protects EsxX-producing cells from self-toxicity prior to EsxX secretion. By contrast ExiCD is encoded by both EsxX producers and in antitoxin islands of competitor strains that do not encode EsxX, consistent with providing immunity against the secreted form of the toxin. This work defines a new class of antibacterial toxin requiring two distinct types of immunity protein which follow different phylogenetic distributions.

## Introduction

*Staphylococcus aureus* is a human nasal commensal and opportunistic pathogen and a major cause of bloodstream infections. Some *S. aureus* strains also colonise animals, in particular cattle and pigs, and are associated with diseases such as mastitis and skin infections^1^. Strain ST398 was originally isolated from swine and identified as a cause of infection for humans working closely with livestock^2,3^. More recently, ST398 clones have been identified that are associated with community-acquired infections in the USA and elsewhere^4,5^.

*S. aureus* encodes a type VII secretion system as part of its core genome. This system, designated the T7SSb, is distantly related to the T7SSa/ESX secretion systems found in mycobacteria. The 5’ end of the *S. aureus ess* locus, which encodes the T7SSb, is highly conserved and codes for the EsxA, EsaA, EssA and EssB components of the secretion machinery (Figure 1A). However, the *ess* locus sequence across strains starts to diverge approximately ¾ of the way through the *essC* gene, which encodes the essential T7SS ATPase component, resulting in four distinct sequence variants, named *essC1* – *essC4*. Downstream of each *essC* variant is a variant-specific set of genes that encode T7SS substrate proteins and accessory factors^6^.

**Figure 1.**
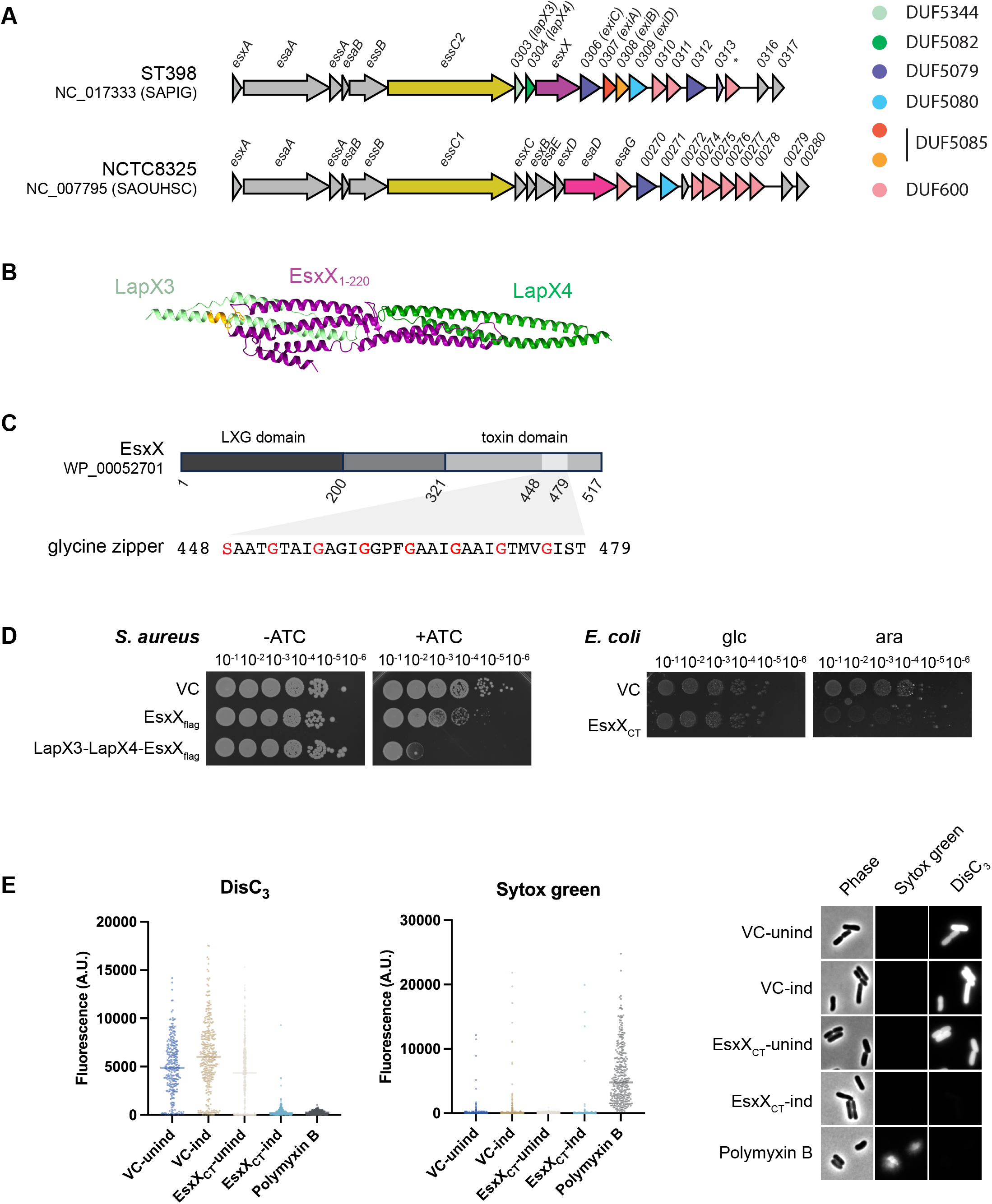
The C-terminal domain of EsxX is a membrane depolarising toxin. A. Schematic diagram of the *S. aureus ess* locus from strains ST398 (top) and NCTC8325 (bottom). Genes are colour-coded by protein family. B. AlphaFold model of LapX3 (SAPIG0303), LapX4 (SAPIG0304) and EsxX_1-220_. The EsxX ‘FxG’ and the LapX3 ‘FxxxD’ targeting motifs are highlighted in orange. Model confidence metrics are shown in figure S1. C. Top: Schematic diagram of the EsxX protein, the glycine zipper motif is shaded light grey. Bottom: Amino acid sequence of the EsxX glycine zipper with motif-defining amino acids in red. D. Left: Overnight cultures of *S. aureus* strain 10.1252.XΔess carrying empty pRab11 (VC) or pRab11 encoding the indicated proteins were serially diluted and spotted onto TSB plates with or without ATC as indicated. Plates were incubated overnight at 37°C. Right: Overnight cultures of *E. coli* strain MG1655 carrying empty pBAD18-cm (VC) or pBAD18-cm encoding the indicated proteins were serially diluted and spotted onto LB plates containing either 1% D-glucose or 0.02% L-arabinose, as indicated. Plates were incubated overnight at 37°C. EsxX_CT_ = residues 321 – 517. E. Exponential phase *E. coli* MG1655 cultures carrying pBAD18-cm (VC) or pBAD18-SAPIG0305-CT (EsxX_CT_) were collected prior to (unind) and post-(ind) induction with 0.2% L-arabinose for 30 minutes. Samples were incubated with the membrane potential-dependent dye DisC_3_(5) and the membrane-impermeant nucleic acid stain Sytox green and imaged by phase contrast and fluorescence microscopy. A control sample carrying pBAD18-cm was treated with polymyxin B for 5 min prior to staining. Data were collected over two independent experiments with n=120-290 cells per condition, single cell fluorescence values are plotted with a line to represent the median. The right hand panel shows representative cell images.

The best characterised T7SS variant is from *essC1* strains. This *essC* variant is found in commonly studied *S. aureus* strains including Newman, RN6390 and USA300. A large nuclease toxin, EsaD is encoded downstream of *essC1*, alongside an immunity protein EsaG, and a globular chaperone EsaE. The nuclease activity of EsaD locates to its C-terminal domain, while an LXG domain is found at the N-terminus^7^. Recently it was shown that three small helical hairpin proteins, EsxB, EsxC and EsxD, also encoded 3’ to *essC1*, interact with the EsaD LXG domain to form a pre-secretion complex. Cryo-EM reveals the complex to comprise a helical shaft formed from the LXG domain and its helical partners and a flexible globular head that presumably encompasses the nuclease domain and the globular proteins EsaE and EsaG^8^.

Two additional T7-secreted antibacterial toxins have since been characterised in strains RN6390 and USA300. TspA is a membrane-depolarising toxin with an N-terminal LXG domain that requires the small helical **<u>L</u>**XG-associated **<u>α</u>**-helical **<u>p</u>**roteins, LapT1 and LapT2, to support its secretion^9,10^. TslA is a phospholipase that has an unusual reverse arrangement of domains; the toxic lipase domain is found at the N-terminus, with a helical LXG-like domain at the C-terminus^11^. TlaA1 and TlaA2 are small helical proteins encoded adjacent to TslA that stack on the helical TslA C-terminus to form a pre-secretion complex. This domain architecture is conserved across other T7SSb toxins, including the *Streptococcus intermedius* LXG-proteins TelA and TelC, each of which also interact with a pair of cognate Lap proteins that are necessary for toxin secretion^12,13^.

To date, the primary role of characterised T7SSb toxins is to mediate interbacterial competition^14^. Each toxin is encoded at a locus that also carries a gene for a cognate immunity protein which serves to protect the producing bacterium. Secretion by the T7SS is post-translational so where antibacterial toxins have a cytoplasmic target, for example EsaD, the producer would be exposed to EsaD toxicity if nuclease activity was not inhibited by EsaG. During secretion it is likely that the globular nuclease domain of EsaD at least partially unfolds, releasing EsaG, which remains in the cytoplasm. Cytoplasmic EsaG also has a second role, which is to protect *S. aureus* from the activity of incoming EsaD secreted by other bacteria. Accordingly, strains of *S. aureus* that lack EsaG proteins are susceptible to EsaD-dependent killing^7^. Toxins that have an extracellular target, for example the lipid II phosphatase, TelC, are also encoded alongside immunity proteins (TipC in the case of TelC) but in this instance the immunity protein is only required to protect from the toxin after it has been secreted, and not during its biosynthesis^15^.

In this study we investigated the biogenesis and activity of the LXG domain protein EsxX, which is encoded at the *essC2* locus of strain ST398. A prior study identified EsxX as a substrate of the T7SSb in this strain, and deletion of the encoding gene resulted in increased bacterial survival and virulence in murine skin and bloodstream infection models^16^. Here we show that EsxX is a membrane-depolarising toxin with an extended glycine zipper motif that has potent antibacterial activity. We demonstrate that EsxX has the unusual property of being able to depolarise the membrane from both the periplasmic and cytoplasmic surface, and that compartment-specific immunity proteins are consequently required to neutralise its activity.

## Results

### EsxX has an N-terminal LXG domain

The *ess* locus in ST398 strains carries a single LXG domain-encoding gene, *esxX*, which was previously identified as a T7SSb substrate^16^ (Figure 1A). Two genes, *SAPIG0303* and *SAPIG0304*, are sandwiched between *essC* and *esxX* and encode small helical proteins of the DUF5344 and DUF5082 families. Proteins of these families are also associated with the LXG toxin TelA in *Streptococcus intermedius* and have been designated LapA3 (DUF5344) and LapA4 (DUF5082) since they were shown to interact with the toxin’s LXG domain^13^. AlphaFold predicts that SAPIG0303 and SAPIG0304 bind the LXG domain of EsxX and we therefore renamed these LapX3 and LapX4, respectively, (<u>L</u>XG-associated <u>α</u>-helical <u>p</u>rotein for Esx<u>X</u>) according to the nomenclature established in *S. intermedius*. LapX3, carries an FxxxD motif on its C-terminal arm, which has been identified as a likely T7SS export signal in other Lap proteins^12^ (Figure 1B, Figure S1A).

### The EsxX C-teminal domain is a toxin with a glycine zipper motif

As an LXG protein, EsxX is predicted to carry a C-terminal toxin domain. While EsxX has no recognisable DUF or PFAM domains apart from the N-terminal LXG domain^17^ (PF04740/IPR006829), BLAST analysis indicated that it shares 22% sequence identity with the T7SS toxin TelE from *Streptococcus gallolyticus* subsp. *gallolyticus* strain UCN34 T7SSb^18^. TelE has been characterised as a membrane-destabilising toxin with an extended glycine zipper motif that is essential for toxic activity against *Escherichia coli*. Glycine zippers are sequence motifs with the pattern (G/A/S)xxxGxxxG or GxxxGxxx(G/S/T), where the repeating GxxxG motif facilitates homo-oligomerisation via close packing of neighbouring transmembrane helices^19,20^. Like TelE, EsxX also harbours an extended glycine zipper motif close to its C-terminus (Figure 1C).

To investigate whether EsxX is toxic to *S. aureus*, we constructed a strain of ST398 that was deleted for the entire T7SS locus (strain 10.1252.XΔess; Table 1; deleted for *esxA* through to *SAPIG0314*) to ensure that any candidate immunity genes were absent. We then expressed a plasmid-borne FLAG-tagged allele of *esxX* from the anhydrotetracycline (ATC)-inducible *P*_*xyl/tet*_ promoter in this strain. Production of cytoplasmic EsxX inhibited growth ten-fold, and co-production of LapX3 and LapX4 further enhanced EsxX toxicity, reflecting a likely role in stabilising EsxX. The EsxX C-terminal domain was also toxic to *Escherichia coli* when produced heterologously from the arabinose-inducible P_BAD_ promoter (Figure 1D).

**Table 1.** *S. aureus* strains.

| Strain | Description | Reference |
| --- | --- | --- |
| 10.1252.X | Livestock-associated ST398 isolate. *essC3* variant strain. | Casabona et al., 2017<sup>45</sup> |
| 10.1252.XΔ*ess* | 10.1252.X with markerless deletion of genes from *SAPIG0297* (*esxA*) to *SAPIG0314*. | This work |

We next probed whether the EsxX C-terminal domain had membrane depolarising and/or pore-forming activity using a single cell fluorescence microscopy approach. *E. coli* producing EsxX_CT_ under control of an arabinose-inducible promoter was incubated with the voltage-sensitive dye DisC3(5). Cells grown in the absence of arabinose or harbouring an empty plasmid vector showed clear fluorescence as the dye accumulated in the membrane due to the presence of a membrane potential. By contrast, cells producing EsxX_CT_ showed barely detectable fluorescence, indicating that they were depolarised (Figure 1E). As depolarisation can result from formation of small ion channels or larger ion-permeable pores, we counter-stained the same cells with the membrane-impermeable nucleic acid stain Sytox green. While *E. coli* cells incubated with Polymyxin B which produces large ion-permeable pores in the *E. coli* cell envelope showed no DisC3(5) fluorescence and strong labelling with Sytox green, EsxX_CT_-producing cells showed no Sytox green staining demonstrating that these cells remained impermeable to larger molecules (Figure 1E). We conclude that EsxX is a membrane depolarising toxin but that, unlike TelE, it does not destabilise the cytoplasmic membrane.

### SAPIG0307 (ExiA) and SAPIG0308 (ExiB) form a dimeric complex for cytoplasmic neutralisation of EsxX

Channel-forming anti-bacterial effectors are often found as part of the type VI secretion system (T6SS) arsenal as well as the T7SSb. In almost all cases, immunity is provided by polytopic membrane proteins^9,21–27^. Two multi-spanning membrane proteins are encoded downstream of EsxX; *SAPIG0306* and *SAPIG0309* each encode a protein with six predicted transmembrane helices, belonging to the DUF5079 and DUF5080 families respectively. However co-production of these proteins together with EsxX did not protect *S. aureus* or *E. coli* from cytoplasmic EsxX toxicity (Figure 2A,B).

**Figure 2.**
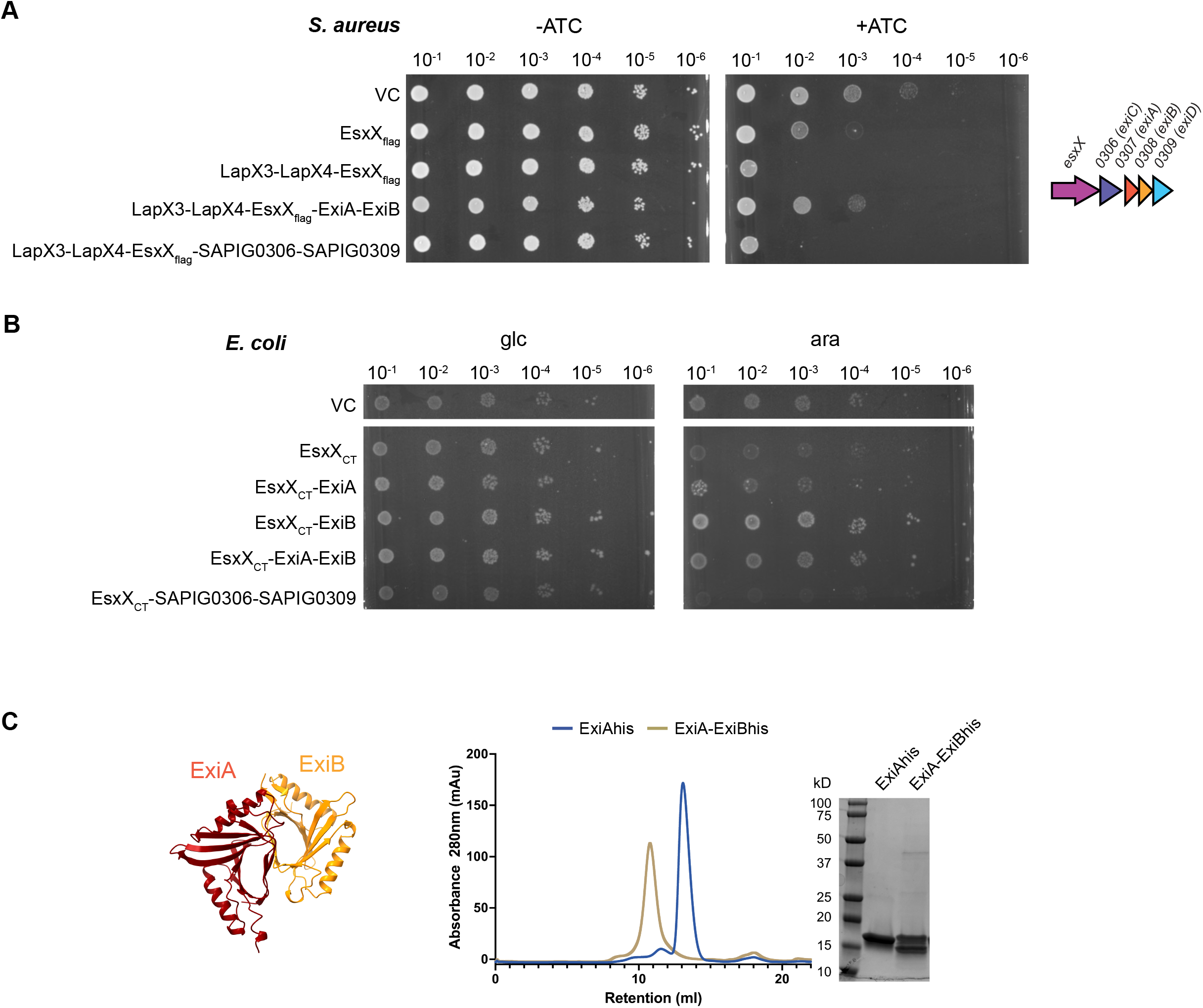
Neutralisation of cytoplasmic EsxX by ExiAB. A. Overnight cultures of *S. aureus* strain 10.1252.XΔess carrying empty pRab11 (VC) or pRab11 encoding the indicated proteins were serially diluted and spotted onto TSB plates with or without ATC as indicated. Plates were incubated overnight at 37°C. B. Overnight cultures of *E. coli* strain MG1655 carrying empty pBAD18-cm (VC) or pBAD18-cm encoding the indicated proteins were serially diluted and spotted onto LB plates containing either 1% D-glucose or 0.02% L-arabinose, as indicated. Plates were incubated overnight at 37°C. EsxX_CT_ = residues 321 – 517. C. ExiAhis and ExiA-ExiBhis were purified from *E. coli* MG1655 cultures carrying pTrc-307h or pTrc-307-308h by nickel affinity chromatography, then analysed by SEC using a Superdex 75 column followed by SDS PAGE with Coomassie staining. On the left is an AlphaFold model of ExiA (SAPIG0307) and ExiB (SAPIG0308). Model confidence metrics are shown in figure S1.

We next examined the remaining genes at the *essC2* locus. The *SAPIG0310* and *SAPIG0311* gene products belong to the DUF600 family which also includes the EsaD immunity protein EsaG. These are therefore likely to be EsaD-neutralising proteins and not EsxX immunity genes. *SAPIG0307* and *SAPIG0308* encode soluble cytoplasmic proteins of the DUF5085 family. AlphaFold predictions suggested that these form a heterodimer, which we confirmed by size exclusion chromatography (SEC) of the purified proteins (Figure 2C, Figure S1), so we tested them as a pair for for their ability to neutralise cytoplasmic EsxX toxicity. As shown in figures 2A and 2B, SAPIG0307-SAPIG0308 provided protection from EsxX toxicity in both *S. aureus* and *E. coli* and we therefore renamed these proteins ExiA and ExiB, respectively (for <u>E</u>sx<u>X</u> Immunity). Production of ExiA or ExiB individually alongside EsxX_CT_ in *E. coli* demonstrated that only ExiB is required for immunity (Figure 1A,B), however parallel expression and purification studies indicated that stable production of EsxX required both ExiA and ExiB (Figure S2). We conclude that soluble proteins mediate protection from the action of EsxX when it is present in the cytoplasm.

### The structural basis for ExiAB-mediated immunity

We next used co-expression and purification in *E. coli* to probe interactions between EsxX and its soluble partners (LapX3, LapX4, ExiA and ExiB). In these experiments an N-terminal his_6_ tag was present on EsxX and a C-terminal twinstrep (ts) tag on ExiB, and complexes were purified via sequential nickel and streptactin affinity chromatography. All five proteins co-purified and when subject to size exclusion chromatography (SEC) were eluted as a single peak (Figure 3A ‘pentamer’). We therefore conclude that all five soluble proteins likely form a stable pre-secretion complex.

**Figure 3.**
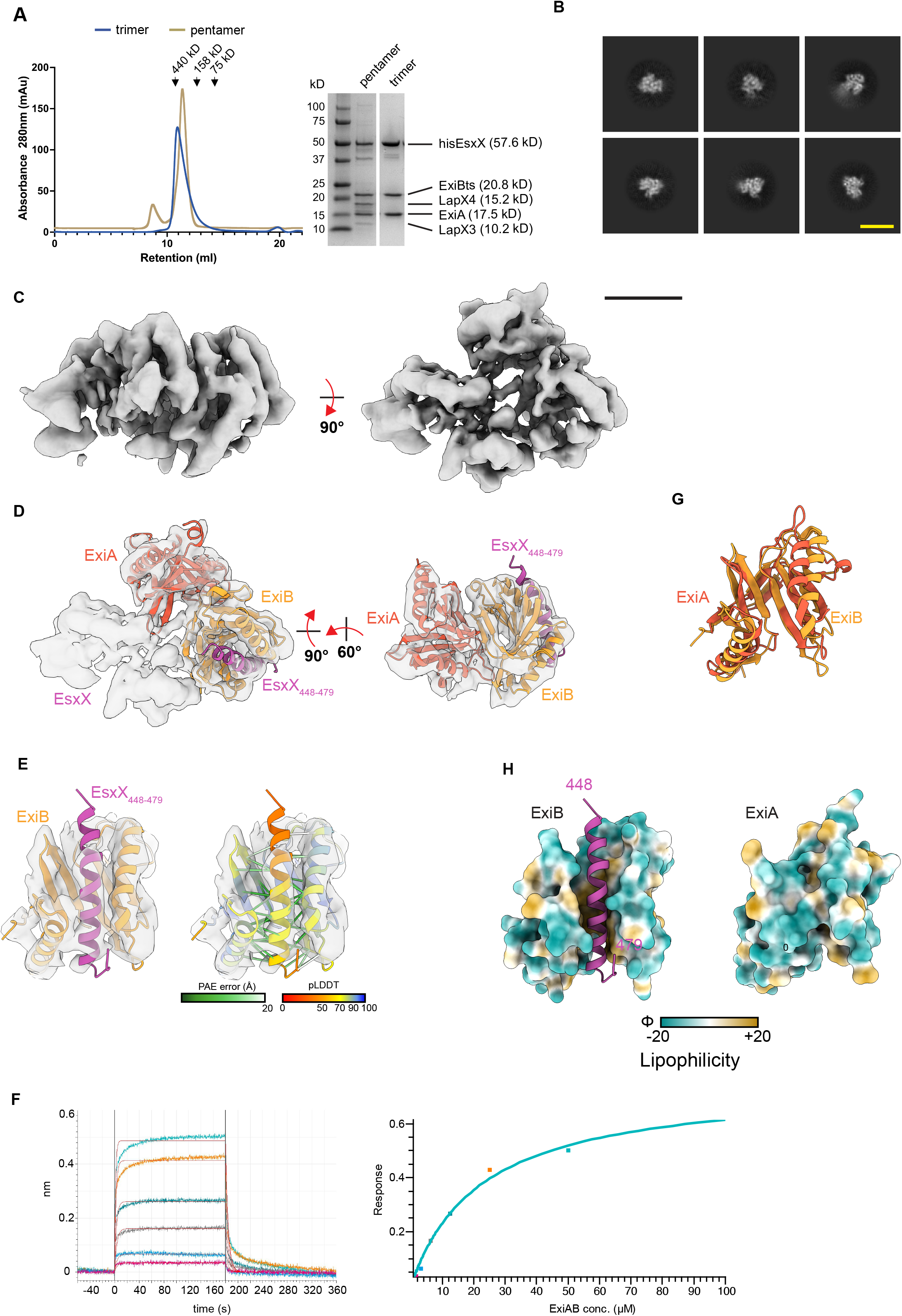
Structural analysis of ExiAB-mediated immunity. A. A hisEsxX-ExiA-ExiBts trimer and a LapX3-LapX4-hisEsxX-ExiA-ExiBts pentamer were purified by sequential nickel and streptactin affinity chromatography, then analysed by SEC using a Superdex 200 increase column, and by SDS PAGE with Coomassie staining. B. Representative cryo-EM 2D class averages of EsxX-ExiA-ExiB. Yellow scale bar, 100 Å. C. Cryo-EM reconstruction of EsxX-ExiA-ExiB, contoured to a threshold of 0.54. Black scale bar, 25 Å. D. AlphaFold model of EsxX_448-479_-ExiA-ExiB fit to cryo-EM density (cross-correlation of 0.6) of the full map (left) or partial map covering EsxX_448-479_-ExiA-ExiB (right). E. Close up view of modelled EsxX_448-479_-ExiB fit into cryo-EM density, coloured by subunit (left) or by pLDDT score (right). EsxX_448-479_-ExiB contacts within 4 Å (Cα-Cα) are displayed as pseudobonds and coloured according to putative alignment error (PAE) score. F. Kinetics (left) and steady state response (right) for ExiAB binding to the EsxX glycine zipper peptide (EsxX_448-476_), assessed by bilayer interferometry. The calculated binding affinity was 22 μM with 1:1 stoichiometry. G. Structural alignment of ExiA and ExiB using the matchmaker function of ChimeraX. Overall Cα rmsd across all residues is 3.9 Å. H. Surface lipophilicity of ExiB with bound EsxX_448-479_ (left) or ExiA (right) in same model orientation as (G).

**Figure 3.** EsxX is toxic when presented to the external face of the membrane, and requires distinct compartment-specific immunity proteins. Overnight cultures of *S. aureus* strain 10.1252.XΔess carrying empty pRab11 (VC) or pRab11 encoding the indicated proteins were serially diluted and spotted onto TSB plates with or without ATC as indicated. Plates were incubated overnight at 37°C. ssEsxX_CT_ denotes the *S. aureus* α-hemolysin signal sequence fused to the C-terminal 203 residues of EsxX.

Based on previous literature together with AlphaFold modelling we expected that binding of LapX3 and LapX4 to the EsxX LXG domain would generate a rod-shaped targeting domain, while the ExiAB immunity complex presumably binds the EsxX toxin domain (Figure S3A). Analysis of this heteropentameric complex by single particle cryo-EM yielded 2D class averages of either rod-like or structured globular complexes (Figure S3B), which likely corresponded to LapX3-LapX4-EsxX and EsxX-ExiA-ExiB subcomplexes, respectively. These results were suggestive of either compositional heterogeneity derived from sample dissociation into subcomplexes upon vitrification or conformational heterogeneity derived from the relative mobility of the subunits, resulting in alignment of either subcomplex but not both together. Either scenario precluded the generation of interpretable cryo-EM maps.

We then focused on reducing compositional and conformational heterogeneity by producing a heterotrimeric complex of only his_6_-EsxX, ExiA and ExiB-ts, again via sequential nickel / streptactin affinity chromatography, followed by SEC (Figure 3A ‘trimer’). This EsxX-ExiA-ExiB complex was imaged by cryo-EM, producing 2D class averages consistent with the globular subcomplex described for the heteropentameric sample, but lacking the rod-like particles as expected (Figure 3B). A moderate resolution cryo-EM volume with dimensions 65 × 75 Å x 55 Å and resolved α-helices was generated for this heterotrimeric complex (Figure 3C). Rigid body fitting the pentameric LapX3-LapX4-EsxX-ExiA-ExiB AlphaFold model into the density showed strong model-map agreement for ExiA-ExiB and the C-terminal glycine zipper of EsxX (residues 448-479), where the glycine zipper is bound by ExiB. A minimal AlphaFold model consisting of EsxX_448-479_-ExiA-ExiB was then generated and rigid-body fitted into the cryo-EM volume (Figure 3D). While the resulting unassigned density likely corresponds to the EsxX toxin domain in a state that AlphaFold could not accurately predict, α-helical density for the EsxX glycine zipper is apparent, showing improved model confidence metrics (pLDDT, PAE) correlating with areas of strong density (Figure 3E). The 1:1 binding of a synthetic EsxX glycine zipper peptide to an ExiAB dimer was confirmed by bilayer interferometry (BLI) (Figure 3F). Both ExiA and ExiB share a compact α/β protein fold with overall dimensions ∼ 45 Å × 35 Å × 30 Å formed from two four-stranded antiparallel β-sheets and two overlying α-helices (Figure 3G). In ExiB these α-helices and a face of the β-sheets form a pronounced hydrophobic groove that recruits the EsxX glycine zipper (Figure 3H). The groove is noticeably absent in ExiA due to packing interactions at the tips of the α-helices, explaining the preferential binding of the EsxX glycine zipper to ExiB. Analysis of 60 ExiB homologues showed a high level of conservation of the hydrophobic residues lining the binding groove (Figure S3C).

### The EsxX C-terminal domain is also toxic when present at the extracellular side of the membrane and is neutralised by ExiC (SAPIG0306) and ExiD (SAPIG0309)

While the T6SS can reportedly deliver toxic effectors directly into the cytoplasm of target cells^28,29^, the T7SS is not a contractile injection system and instead delivers toxins into the surroundings or to the surface of competitor bacteria. Therefore, EsxX would be expected to encounter the cytoplasmic membranes of bacterial targets from the extracellular side. To investigate whether the EsxX toxin domain was also toxic when present outside of the cytoplasm, we appended a Sec signal peptide to the N-terminus of EsxX_CT_ and produced this in *S. aureus* and found that the periplasmically-targeted EsxX_CT_ was highly toxic (Figure 4A). We conclude that the C-terminal 196 amino acids of EsxX is an antibacterial toxin which is active in both the cytoplasm and periplasm of bacteria.

**Figure 4.**
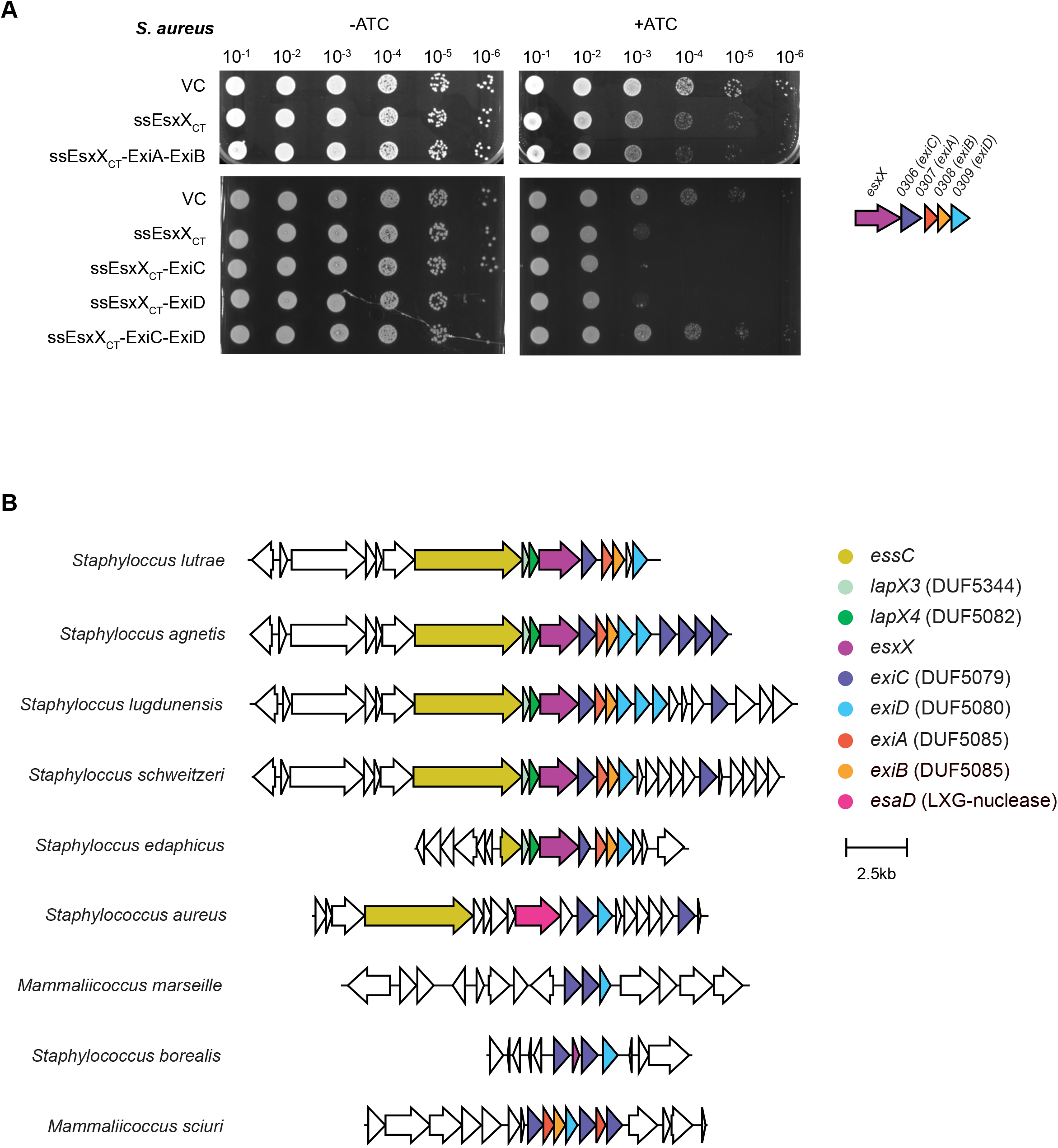
Genetic loci of homologues of EsxX and ExiC. Genetic loci of staphylococcal *esxX* (magenta) and its immunity genes, identified by FlaGs analysis of EsxX, ExiC (SAPIG0306) and ExiB (SAPIG0308) (complete data are shown in figures S4-S6). Homologous genes are colour-coded. The strains depicted are as follows, with genome accessions given in brackets: *S. lutrae* ATCC700373 (NZ_CP020773), *S. agnetis* 1379 (NZ_CP045927), *S. lugdunensis* FDAARGOS_141 (NZ_CP014022), *S. schweitzeri* NCTC13712 (NZ_LR134304.1), *S. edaphicus* CCM 8730 (NZ_MRZN01000025.1), *S. aureus* RN4220 (NZ_CP101124), *M. Marseille* Q6498 (NZ_OX267714), *S. borealis* 58-52 (NZ_JABVEF010000003), *M. sciuri* SNUC 1345 (NZ_QYJC01000003).

As ExiAB were able to neutralise cytoplasmic EsxX_CT_ toxicity we tested whether they could also provide protection against the toxin when it was present extracellularly (Figure 4A). However, in this instance the immunity proteins did not alleviate toxicity, consistent with them acting only in the cytoplasmic compartment. We next tested whether either of the two polytopic membrane proteins, SAPIG0306 and SAPIG0309 could protect against EsxX_CT_ extracellular toxicity. While neither protein inhibited toxicity when produced alone, co-producing both of them offered complete protection against the action of extracellular EsxX_CT_. In agreement with a requirement for both of these to neutralise secreted EsxX, AlphaFold predicted the two proteins to form a heterodimer (Fig. S1C). These proteins were therefore renamed ExiC (SAPIG0306) and ExiD (SAPIG0309).

### *exiC* and *exiD* occur as orphan immunity genes in staphylococcal genomes

To explore the distribution of *esxX* and its immunity genes across staphylococcaceae we undertook gene neighbourhood analysis. As expected, an *exiAB* gene pair is located downstream of all *esxX* genes analysed, consistent with the requirement for ExiAB to protect the producing bacteria from cytoplasmic EsxX toxicity prior to secretion by the T7SS (Figures 4B, S4, S5). By contrast, *exiCD* were not only encoded at *esxX* loci, but were also found in many genomes that did not encode *esxX*/*exiA*/*exiB*, or even T7SS structural genes (e.g. *Staphylococcus borealis*). In these instances, the *exiC* and *exiD* occurred together, located in ‘antitoxin islands’^30^ (Figure 4B, Figure S6). These are regions of staphylococcal genomes that are enriched in multiple T7SS immunity genes that are likely acquired due to selective pressure of T7SS toxin-mediated bacterial antagonism. Taken together, these findings strongly suggest that a major biological role of EsxX is to mediate intra-/inter-species competition.

## Discussion

In this work we have characterised the EsxX toxin of the *S. aureus essC2* variant strain ST398. We show that the EsxX C-terminal domain has membrane-depolarising activity that is toxic to bacteria. Our data support a model where secreted EsxX antagonises competitor bacteria by depolarising the cytoplasmic membrane of susceptible cells, likely mediated by membrane insertion of the EsxX glycine zipper and assembly into ion channels, as described for other glycine zipper toxins^31^. Toxicity is prevented by a membrane-integrated pair of proteins, ExiC and ExiD. In agreement with ExiCD acting as immunity factors to protect from secreted EsxX toxicity, genes encoding these proteins are frequently found as orphans in genomes of bacteria that do not code for *esxX*, clustering in genomic antitoxin islands with other T7SS immunity genes.

Surprisingly, EsxX also exhibits toxicity when present in the cytoplasm, necessitating the requirement for the producing strains to encode an additional immunity protein family. ExiA and ExiB are a pair of DUF5085 proteins and form a heterodimer which binds to the glycine zipper region of cytoplasmic EsxX to prevent toxicity. Phyre2 predictions reveal that ExiA and ExiB are structurally homologous to proteins of the GyrI-like superfamily (IPR029442), whose members carry a duplicated SHS2 (strand-helix-strand-strand) fold (Figure S7). This fold is usually adapted for promiscuous binding of small molecules^32,33^, for example the *Bacillus subtilis* BmrR transcription factor, which functions in multidrug resistance, is activated by binding of diverse ligands to its GyrI-like effector binding domain^34–36^. Here the same binding cleft in ExiB, between the two helices of the two SHS2 domains, is adapted for protein-protein interactions and is lined with hydrophobic residues. It should be noted that TipE, the second of two DUF5085 proteins encoded downstream of *telE* was shown to neutralise TelE toxicity in *Streptococcus gallolyticus*^18^, in agreement with our findings that ExiB directly binds the toxin’s glycine zipper region.

The *S. aureus* T7SS secretes a second membrane-depolarising toxin, TspA. TspA has a shorter glycine zipper motif than EsxX and only exhibits toxicity from the extracellular side of the membrane. Accordingly, *S. aureus* encodes a single membrane-bound immunity protein TsaI for protection from TspA toxicity following secretion^9^. Membrane-depolarising toxins are also secreted by the Gram-negative T6SS, for example Tse5 of *Pseudomonas aeruginosa* whose toxin domain is found at the C-terminus of a large bacterial rearrangement hotspot (Rhs) protein. Like EsxX, the C-terminus of Tse5 exhibits both cytoplasmic and periplasmic toxicity^22,23,37^. Tsi5 is a membrane-bound immunity protein that protects cells from extracellular Tse5^22,23^, however there is no requirement for a cytoplasmic immunity protein because the Rhs repeats of Tse5 form a barrel-like structure with a plug domain that encapsulates the toxic C-terminus, shielding it from activity in the cytoplasm^37^.

We propose a model for the biogenesis of EsxX, illustrated in figure 5. The cytoplasmic form of EsxX in *S. aureus* ST398 exists as a pre-secretion complex where LapX3 and LapX4 bind to the N-terminal LXG domain and ExiAB to the toxin domain, sequestering the glycine zipper. A composite signal sequence formed from the LXG domain and Lap proteins targets the complex to the T7SSb. During secretion the hexameric EssC secretion channel is predicted to open to a diameter of ∼30 Å^38^, accommodating the folded LXG rod domain and Lap partners but not the more bulky ExiAB complex. This would result in stripping of the immunity proteins from the unfolded toxin domain and release of the active toxin into the environment. The toxin may then access the envelope of target cells to insert into and depolarise the membrane unless the strain expresses the ExiCD membrane-bound immunity proteins.

**Figure 5.**
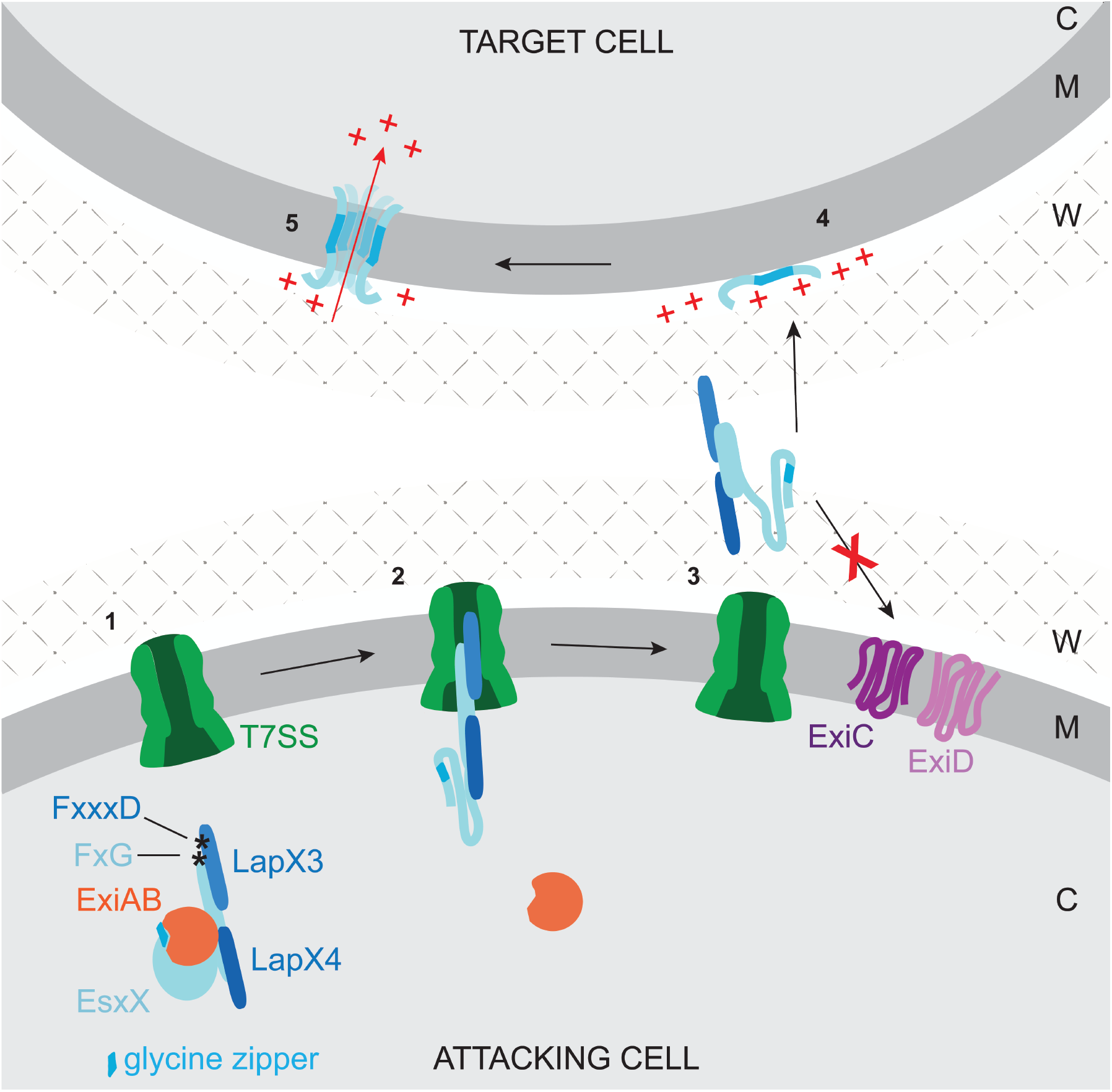
Model for EsxX biogenesis. 1. The ExiAB immunity complex binds to the toxin domain of nascent EsxX in the cytoplasm, preventing its membrane insertion. LapX3 and LapX4 assemble with the EsxX LXG domain, forming a rod-shaped transport complex carrying a composite FxG-FxxxD targeting signal. 2. The FxG-FxxxD signal delivers the EsxX presecretion complex to the T7SS apparatus at the cytoplasmic membrane. As the rod-shaped complex traverses the secretion channel, ExiAB is stripped from the toxin domain, which is carried through the channel in an unfolded conformation. 3. Following toxin secretion, ExiC and ExiD protect the attacking cell membrane from insertion and/or assembly of the EsxX glycine zipper 4. The LXG domain and/or Lap proteins might be removed from the toxin, which then targets the competitor cell membrane. 5. The toxin inserts into the target membrane and oligomerises via its glycine zipper, causing depolarisation of the target cell membrane. C: cytoplasm, M: membrane, W: cell wall, +: positively charged ion

## Methods

### Bacterial strains, plasmids and growth conditions

*E. coli* was cultured in Luria broth (LB, Melford) or terrific broth (TB, Formedium) with ampicillin (100 μg/ml) and/or chloramphenicol (25 μg/ml) where required. *S. aureus* was cultured in tryptic soy broth (TSB; Oxoid) with chloramphenicol (10 μg/ml) where required. *E. coli* strains DH5???????? and JM110 were used for cloning and preparation of plasmids for electroporation respectively. *E. coli* MG1655 (F-lambda-*ilvG*-*rfb*-50 *rph*-1) was used to assay toxicity of EsxX_CT_ and for fluorescence microscopy to assess changes to membrane potential and permeability. *E. coli* MG1655 and BL21(DE3) (F^−^ *omp*T *hsd*S_B_ (r_B–_, m_B–_) *gal dcm* (DE3)) were used for overproduction and purification of recombinant proteins. *S. aureus* strains and all plasmids used are listed in Tables 1, 2 and S1. Details of how strains and plasmids were constructed are provided as supplementary information.

**Table 2.** Plasmids.

| Plasmid | Details | Ref/source |
| --- | --- | --- |
| pRab11 | *E. coli*/*S. aureus* shuttle vector, carries P<sub>xyl/tet</sub> for inducible expression. Amp<sup>r</sup>, cml<sup>r</sup>. | Helle et al., 2011<sup>46</sup> |
| pRab11-SAPIG0305-flag | pRab11 producing full length EsxX with a C-terminal FLAG tag | This work |
| pRab11-303-304-305flag | pRab11 producing LapX3, LapX4 and EsxX with a C-terminal FLAG tag. | This work |
| pRab11-303-304-305flag-307-308 | pRab11 producing LapX3, LapX4, EsxX-FLAG, ExiA and ExiB. | This work |
| pRab11-303-304-305flag-307 | pRab11 producing LapX3, LapX4, EsxX-FLAG and ExiA. | This work |
| pRab11-303-304-305flag-308 | pRab11 producing LapX3, LapX4, EsxX-FLAG and ExiB. | This work |
| pRab11-303-304-305flag-306-309 | pRab11 producing LapX3, LapX4, EsxX-FLAG, ExiC and ExiD. | This work |
| pRab11-Hlass-SAPIG0305CT | pRab11 producing ssHla-EsxX<sub>CT</sub> (*S. aureus* Hla signal sequence fused to amino acids 317-517 of SAPIG0305). | This work |
| pRab11-Hlass-305CT-306-309 | pRab11 producing ssHla-EsxX<sub>CT</sub>, ExiC and ExiD. | This work |
| pRab11-Hlass-305CT-306 | pRab11 producing ssHla-EsxX<sub>CT</sub> and ExiC. | This work |
| pRab11-Hlass-305CT-309 | pRab11 producing ssHla-EsxX<sub>CT</sub> and ExiD. | This work |
| pRab11-Hlass-305CT-307-308 | pRab11 producing ssHla-EsxX<sub>CT</sub>, ExiA and ExiB | This work |
| pBAD18-cm | Glucose-repressible/arabinose-inducible vector; cml<sup>r</sup>. | Ulhuq et al., 2020<sup>9</sup> |
| pBAD18-SAPIG0305-CT | pBAD18-cm producing EsxX<sub>CT</sub> (residues 321-517) | This work |
| pBAD305CT-307-308 | pBAD18-cm producing EsxX<sub>CT</sub> (residues 321-517), ExiA and ExiB | This work |
| pBAD305CT-307 | pBAD18-cm producing EsxX<sub>CT</sub> (residues 321-517) and ExiA | This work |
| pBAD305CT-308 | pBAD18-cm producing EsxX<sub>CT</sub> (residues 321-517) and ExiB | This work |
| pBAD305CT-306-309 | pBAD18-cm producing EsxX<sub>CT</sub> (residues 321-517), ExiC and ExiD | This work |
| pTrc99A | Bacterial expression vector with inducible lacI promoter; amp<sup>r</sup> | Amann et al., 1988 Gene<sup>47</sup> |
| pTrc-307h | pTrc99A producing ExiA with a C-terminal his<sub>6</sub> tag. | This work |
| pTrc-307-308h | pTrc99A producing ExiA and ExiB with a C-terminal his<sub>6</sub> tag. | This work |
| pTrc-307-308ts | pTrc99A producing ExiA and ExiB with a C-terminal twinstrep tag. | This work |
| pACYCDuet-1 | Bacterial expression vector with inducible T7 promoter and lac operator; cml<sup>r</sup>. | Novagen |
| pACD-his305 | pACYCDuet producing EsxX with an N-terminal his<sub>6</sub> tag. | This work |
| pTrc-303-304-h305-306-307-308ts | pTrc99A producing LapX3, LapX4, EsxX with an N-terminal his<sub>6</sub> tag, SAPIG0306, ExiA and ExiB with a C-terminal twinstrep tag. | This work |
| pIMAY-ess-ST398 | pIMAY carrying the regions upstream of esxA and downstream of SAPIG0314, for construction of 10.1252.XΔess | This work |

### Fluorescence microscopy

Cells were assessed for changes in membrane potential and permeabilization as previously described^9^. Briefly, *E. coli* MG1655 cells harbouring pBAD18-Cm derivatives were cultured aerobically in LB. Uninduced and arabinose-induced samples were collected and stained with 2 μM DiSC_3_ and 200 nM Sytox Green (both from Thermofisher) and immobilised on microscope slides in 1.2 % agarose. Fluorescence and phase contrast images were collected using a Nikon Eclipse Ti and analysed using ImageJ/Fiji. A minimum of 280 cell images, collected over two independent experiments, were analysed for each condition.

### Protein purification

For purification of ExiA, the ExiA-ExiB dimer, and the LapX3-LapX4-EsxX-ExiA-ExiB pentamer, cultures of MG1655 cells carrying pTrc-307h, pTrc-307-308h or pTrc-303-304-h305-306-307-308ts were diluted 1/40 into LB containing ampicillin (100 μg/ml) and cultured for 2 h at 37°C. Following addition of 0.1 mM IPTG (isopropyl β-d-1-thiogalactopyranoside) cultures were grown overnight at 22°C. Cells were harvested by centrifugation (15 min, 4000 g, 4°C) and resuspended in Buffer A (50 mM Tris.Cl pH 7.4, 150 mM NaCl) containing 0.2 mg/ml lysozyme, before sonication to disrupt the cells. After centrifugation of the cell lysate (30 min, 25000 g, 4°C), 5 mM (for the monomor and dimer) or 20 mM (for the pentamer) imidazole was added to the supernatant which was subsequently applied to a HisTrap FF column (Cytiva) and washed extensively with Buffer A + 5/20 mM imidazole. Bound proteins were eluted with a gradient up to 500 mM imidazole. The monomer/dimer were then applied to a Superdex 75 10/300 GL column (Cytiva) pre-equilibrated in Buffer A. For the pentamer, protein eluted from the HisTrap column was applied to a Strep-tactin XT column (IBA, 2-4021-001), washed with buffer W (100 mM Tris-HCl, pH 8.0, 150 mM NaCl, 1 mM EDTA) then eluted in buffer BXT (IBA 1042-025). Eluted protein was concentrated using a 50 MWCO centrifugal concentrator and injected onto a Superdex 200 increase 10/300 column (Cytiva) pre-equilibrated in buffer C (50 mM Hepes pH 7.5, 150 mM NaCl).

For purification of an EsxX-ExiA-ExiB trimer, BL21(DE3) cells carrying pACDhis305 and pTrc307-308ts were cultured overnight in LB broth supplemented with chloramphenicol (25 μg/ml) and ampicillin (100 μg/ml). Overnight cultures were diluted 1/50 into TB containing the same antibiotics, then cultured for 1h45 at 37°C, before induction with 50 μM IPTG and overnight growth at 20°C. Cell pellets were resuspended in buffer B (50 mM Tris.Cl pH 7.4, 300 mM NaCl) containing lysozyme (0.2 mg/ml), PMSF (1 mM) and complete mini protease inhibitor cocktail (Roche). Cells were then lysed by sonication, and clarified by centrifugation for 30 min at 25000 g, 4°C. The resulting supernatant was applied to a Histrap FF column (Cytiva) and washed in buffer B containing 20 mM imidazole, then eluted using a 20-250 mM imidazole gradient. Fractions containing EsxX, ExiA and ExiB were exchanged into buffer W using HiTrap desalting columns (Cytiva) then applied to a Strep-tactin XT column (IBA, 2-4021-001), washed with buffer W (100 mM Tris-HCl, pH 8.0, 150 mM NaCl, 1 mM EDTA) then eluted in buffer BXT (IBA 1042-025). Eluted protein was exchanged into buffer C using a HiTrap desalting column.

### Cryo-EM sample preparation, imaging and data processing

LapX3-LapX4-EsxX-ExiA-ExiB (0.3 mg/ml) and EsxX-ExiA-ExiB (0.4 mg/ml) samples were adsorbed onto glow-discharged holey carbon-coated grids (Quantifoil 300 mesh, Au R1.2/1.3) for 10 s. Grids were then blotted for 3 s at 10 °C, 100% humidity and frozen in liquid ethane using a Vitrobot Mark IV (Thermo Fisher Scientific). For EsxX-ExiA-ExiB trimer, 0.15 mM of fluorinated octyl maltoside (Anatrace) was added to the sample prior to adsorption on the grid.

Grids of LapX3-LapX4-EsxX-ExiA-ExiB were collected on a Talos Arctica (Thermo Fisher Scientific) operating at 200 kV, with a Bioquantum GIF (Gatan) with slit width set to 20 eV, at 100,000x magnification with a calibrated pixel of 0.81 Å on a K3 direct electron detector (Gatan). 1,111 movies were collected, each with a total dose of 44.5 e-/A^2^, fractionated to ∼ 1 e™ / Å^2^ / fraction for motion correction. Grids of EsxX-ExiA-ExiB trimer were collected in Electron Event Representation (EER) format, on a CFEG-equipped Titan Krios G4 (Thermo Fisher Scientific) operating at 300 kV with a Selectris X imaging filter (Thermo Fisher Scientific) with slit width of 10 eV at 165,000x magnification on a Falcon 4i direct detection camera (Thermo Fisher Scientific) corresponding to a calibrated pixel size of 0.732 Å. 6,170 movies were collected at a total dose of 56.7 e-/Å2, fractionated to ∼ 1 e™ / Å2 / fraction for motion correction.

Movie preprocessing (patched motion correction, CTF estimation, particle picking and extraction) and initial 2D classification was performed in SIMPLE 3.0^39^. For LapX3-LapX4-EsxX-ExiA-ExiB particles, two rounds of 2D classification was performed in cryoSPARC^40^ using a soft circular mask of 120 Å. 154,103 particles were recovered after 2D classification and subjected to multi-class (k=3) *ab initio* volume generation followed by non-uniform refinement, but the volumes generated were low resolution (6-7 Å) which deterred modeling. For EsxX-ExiA-ExiB trimer two rounds of 2D classification was performed in cryoSPARC using a soft circular mask of 140 Å. This resulted in the selection of 349,850 pruned particles which were used as input in multi-class (k=4) *ab initio* reconstructions. Particles (105,726) from the most prominent class were selected and non-uniform refined against their corresponding volume lowpass-filtered to 15 Å. This generated a 3.7 Å volume, as estimated from gold-standard Fourier shell correlations (FSCs) using the 0.143 criterion, but suffered from significant resolution anisotropy (cFAR score of 0.03) which could not be alleviated by further 2D/3D classification schemes. ChimeraX (pettersen 2021) was used to generate figure panels of maps and models.

### BLI

Binding of purified ExiAB to the EsxX glycine zipper peptide was measured using an Octet^®^ R8 biolayer interferometry (BLI) system (Octet^®^, Sartorius). A peptide encompassing residues 448-476 of EsxX was synthesised with an N-terminal biotin label (biotin-SAATGTAIGAGIGGPFGAAIGAAIGTMVG) by Severn Biotech. The biotinylated peptide was prepared to a final concentration of 250 µg/mL in 10X Kinetics Buffer (KB, (Octet^®^, Sartorius)) containing a final concentration of 1% DMSO. Streptavidin-coated Octet^®^ SAX biosensors (Octet^®^, Sartorius) were hydrated in 10X KB for 10 minutes prior to capture of the biotinylated peptide to a final response level of ∼ 1.0 nm after baseline stabilisation in 10X KB. Purified ExiABhis was diluted to a concentration of 50 μM in 10X KB, then serial two-fold dilutions were made down to 1.6 μM in the same buffer. Association between ExiABhis and the peptide was measured over 180 s at 25 °C. Data were double reference subtracted against the buffer and buffer-loaded biosensors, and fitted to both a 1:1 kinetic model and a steady state binding model using the average response over the time period from 170 – 175 seconds.

### Miscellaneous methods

Gene neighbourhood analysis was carried out using the WebFlaGs server^41^. Figures showing gene clusters were created using Clinker^42^. Protein structures, maps and models were illustrated using ChimeraX^43^ and AlphaFold PAE charts in figure S1 were downloaded from PAE viewer^44^.

## Supporting information

EsxX SI

## Acknowledgements

This study was supported by the Wellcome Trust (through Investigator Awards 10183/Z/15/Z and 224151/Z/21/Z to TP), the Intramural Research Program of the NIH, NCI, Center for Cancer Research (awarded to SL), the China Scholarship Council (through award of a PhD studentship to YY) and the University of Newcastle. We thank Stuart Knowling and Sartorius for providing materials and assistance with the BLI experiment.

## Author contributions

Conceptualization (TP, GC, FA); Investigation (FA, YY, JD, GC, CM, FU); Writing – original draft (FA, JD, TP) and funding acquisition (TP and SL).

