## Supplementary material for "Distinct immunity protein families mediate compartment-specific neutralisation of a bacterial toxin": EsxX SI

### Supplementary Information

[1] Newcastle University Biosciences Institute, Newcastle University, Newcastle upon Tyne, NE2 4HH, UK.

[2] Center for Structural Biology, Center for Cancer Research, National Cancer Institute, NIH, Frederick, MD 21702.

[†] contributed equally

***SI Figure ends***

**Figure S1.**

AlphaFold multimer models for (a) LapX3 (SAPIG0303) and LapX4 (SAPIG0304) in complex with the EsxX (SAPIG0305) LXG domain (residues 1-220), (b) ExiA (SAPIG0307) with ExiB (SAPIG0308), and (c) ExiC (SAPIG0306) with ExiD (SAPIG0309). The EsxX 'FxG' and the LapX3 'FxxxD' targeting motifs in (a) are highlighted in orange. Each model is shown in duplicate, coloured by subunit or by pLDDT score, and the predicted aligned error (PAE) plot is shown below each model. 'EC' – extracellular; 'cyto' – cytoplasm.

**Figure S2**

*E. coli* BL21(DE3) cultures carrying pACD-his305 for production of hisEsxX, together with a pTrc99-derivative plasmid producing the indicated ExiA / ExiB protein(s) were induced with IPTG. Whole cell lysates were analysed by immunoblot with antibodies against his or strep tags. EsxX was only detected in the presence of both ExiA and ExiB. * denotes a degradation product.

**Figure S3**

A. AlphaFold model of a LapX3-LapX4-EsxX-ExiA-ExiB pentamer coloured by subunit (i) or by pLDDT score (ii), with corresponding PAE plot.

B. Representative cryo-EM 2D class averages of purified LapX3-LapX4-EsxX-ExiA-ExiB pentamer. Yellow scale bar, 100 Å.

C. Conservation analysis of 60 DUF5085 sequences identified by BLAST as homologues of the glycine zipper immunity proteins ExiB (SAPIG0308), SAR0290 or TipE (GALLO_0565).

**Figure S4**

Flanking genes (FlaGs) analysis of staphylococcal EsxX (SAPIG0305) homologues which are aligned and shaded black. Genes 1, 6, 3 and 2 encode proteins belonging to the DUF5079, DUF5085, DUF5085 and DUF5080 families respectively.

**Figure S5**

FlaGs analysis of ExiB (SAPIG0308) homologues, which are aligned and shaded black. Gene 5 corresponds to *esxX.*

**Figure S6**

FlaGs analysis of homologues of the DUF5079 protein ExiC (SAPIG0306), which are aligned and shaded black or orange. Genes numbered 2 and 3 correspond to additional DUF5079- and DUF5080-encoding genes respectively. Genes numbered 1 encode homologues of the nuclease immunity protein EsaG. Genes numbered 8 are *esxX* homologues, and gene pairs numbered 5-6 are *exiAB* homologues.

**Figure S7**

Comparison of the ExiA-ExiB AlphaFold model and *E. coli* GyrI crystal structure (1JYH)[1]. Images on the right hand side are rotated 90° to show the binding groove between the two helices. For clarity, the ExiA structure is not shown in the rotated view.

### *SI Methods*

#### Construction of strains and plasmids

Oligonucleotides and templates used for PCR are listed in Table S1. Plasmid deletions were achieved using the Q5 site directed mutagenesis kit (NEB).

Chromosomal deletion of the *ess* locus in 10.1252.X was accomplished by allelic exchange using plasmid pIMAY[2] carrying the flanking regions (~500 bp each) of the deleted region. pIMAY-ess-ST398 was created by HiFi assembly (NEB).

pBAD18-SAPIG0305-CT for *E. coli* toxicity assays was created by PCR-based restriction cloning using the NheI and SalI sites of pBAD18-cm. A *SAPG0307-SAPIG0308* fragment was inserted downstream of SAPIG0305-CT in pBAD18-SAPIG0305-CT by restriction cloning (SalI-SphI) to make pBAD305CT-307-308. *SAPIG0307* (*exiA*) and *SAPGI0308* (*exiB*) were then individually deleted from this plasmid to create pBAD305CT-308 and pBAD305CT-307 respectively. For co-production of SAPIG0306 and SAPIG0309 with $EsxX_{CT}$, *SAPIG0306* was inserted downstream of *SAPIG0305CT* by restriction cloning (SalI-SphI), then *SAPIG0309* was inserted downstream of *SAPIG0306* using HiFi assembly.

pRab11-SAPIG0305-flag was constructed by PCR-based restriction cloning using the KpnI and SacI sites of pRab11. pRab11-303-304-305flag was constructed in multiple steps: First a fragment encompassing *SAPIG0307* to *SAPIG0309* was cloned into pRab11 using PCR and restriction digestion, then a fragment from *SAPIG0303* to *SAPIG0306* was inserted upstream of this using HiFi assembly. The C-terminal FLAG tag on *SAPIG0305* was then inserted (creating pRab11-303-304-305flag-306-307-308-309) and genes from *SAPIG0306* to *SAPIG0309* were deleted (creating pRab11-303-304-305flag), both using the Q5 site-directed mutagenesis kit NEB). For pRab11-303-304-305flag-306-309, *SAPIG0307-SAPIG0308* was deleted from pRab11-303-304-305flag-306-307-308-309 described above. For pRab11-303-304-305flag-307-308, *SAPIG0307-SAPIG0308* was inserted into pRab11-303-304-305flag by HiFi assembly. For pRab11-303-304-305flag-307 and pRab11-303-304-305flag-308, inserts encompassing the required genes were amplified from pRab11-303-304-305flag-307-308 and ligated with the pRab11 backbone by HiFi assembly. For pRab11-Hlass-SAPIG0305CT, fragments corresponding to the Hla signal sequence and $EsxX_{CT}$ were amplified individually then joined together by overlap PCR, then the whole Hlass-SAPIG0305CT fragment was cloned KpnI-SacI into pRab11. *SAPIG0306* and *SAPIG0309* were subsequently cloned downstream of Hlass-SAPIG0305CT by HiFi assembly, creating pRab11-Hlass-305CT-306-309, pRab11-Hlass-305CT-306 and pRab11-Hlass-305CT-309.

For pTrc-307h, *SAPIG0307* was amplified with a C-terminal his tag and cloned NcoI-HindIII into pTrc99A. For pTrc-307-308h, a *SAPIG0307-SAPIG0308* fragment was amplified with a C-terminal his tag on *SAPIG0308*, and cloned NcoI-HindIII into pTrc99A. This plasmid was modified to encode a twinstrep tag in place of the his tag using the Q5 site-directed mutagenesis kit (NEB) to make pTrc-307-308ts. To make pTrc303-304-h305-306-307-308ts a fragment encompassing genes from *SAPIG0303* to *SAPIG0306* was inserted into pTrc-307-308ts by HiFi assembly, then a his tag was inserted at the *SAPIG0305* (*esxX*) N-terminus using the Q5 site-directed mutagenesis kit. For pACD-his305, *SAPIG0305* was cloned BamHI-SalI into pACYCDuet-1 MCS1 for expression with an N-terminal his tag.

**Miscellaneous**

Anti-his antibodies were from Invitrogen (11533923) and anti-strep antibodies were from Qiagen (34850).

Table S1. Oligonucleotides

| Name | Sequence | Template | Construct |
|---|---|---|---|
| MGC414 | tatcgataagcttgatatcgGTGAATCAATTGCGTCATTAATG | 10.1252.X gDNA | pIMAY-ess-ST398 |
| MGC415 | ttcatcacctTAATCATTGCCATAACTAGAAAC | 10.1252.X gDNA | pIMAY-ess-ST398 |
| MGC416 | gcaatgattaAGGTGATGAATTAAATTATTATTCAG | 10.1252.X gDNA | pIMAY-ess-ST398 |
| MGC417 | tggatcccccgggctgcaggAACATTTTAATAGTATATACTTTTCTCATATTTTTTTG | 10.1252.X gDNA | pIMAY-ess-ST398 |
| 2fwd305Nhe1C-term | GCGCGCTAGCCAGAGGAGGAGCCATGAAAATGAAATCCAGTGAGAAATTAAAAGC | 10.1252.X gDNA | pBAD18-SAPIG0305-CT |
| 2rvs305_stop_SalI | GCGCGTCGACTTAAAATACATTGCTTAACGTTTT | 10.1252.X gDNA | pBAD18-SAPIG0305-CT |
| FA221_SalI_307us_F | GGA**GTCGAC**TTGGAGGAATAATAAATGGAAC | 10.1252.X gDNA | pBAD305CT-307-308 |
| FA245_SphI_308_R | GGA**GCATGC**TCAATATTCCATAACTTTTACTTTAATATC | 10.1252.X gDNA | pBAD305CT-307-308 |
| FA249_D308_Q5R | TCATTCCAATTCATCCTCATCTAAATTG | pBAD305CT-307-308 | pBAD305CT-307 |

| | | | |
|---|---|---|---|
| FA247_D308_Q5F | GCATGCAAGCTTGGCTGT | pBAD305CT-307-308 | pBAD305CT-307 |
| FA250_D307_Q5F | TGAGGATGAATTGGAATG | pBAD305CT-307-308 | pBAD305CT-308 |
| FA251_D307_Q5R | GTCGACTTAAAATACATTGC | pBAD305CT-307-308 | pBAD305CT-308 |
| MGC210 | GCGCGTCGACCAGAGGAGGAGCCATGAATAATACTAAGG | 10.1252.X gDNA | pBAD305CT-306-309 |
| MGC211 | GCGCGCATGCTTAGTCATCATCTGCTGTG | 10.1252.X gDNA | pBAD305CT-306-309 |
| FA_pBADssPB305CT_306_fwd | AAGCTTGGCTGTTTTGGC | pBAD305CT-306 | pBAD305CT-306-309 |
| FA_pBADssPB305CT_306_rev | GCATGCTTAGTCATCATCTG | pBAD305CT-306 | pBAD305CT-306-309 |
| FA_rbs309 plus ds (pTrc)_fwd | cagatgatgactaagcatgcGGAATATTGAGGTGCGAATATGGAGTTC | 10.1252.X gDNA | pBAD305CT-306-309 |
| FA_rbs309 plus ds (pTrc)_rev | ccgccaaaacagccaagcttTGCATGCCTGCAGGTCGA | 10.1252.X gDNA | pBAD305CT-306-309 |
| for_KpnI_SAPIG0305 | GCGCGGTACCAGGAGGTTTCTAGTTATGGGGAATAAAATAAAAATGTC | 10.1252.X gDNA | pRab11-SAPIG0305-flag |
| rev_SacI_flag_SAPIG0305 | GCGCGAGCTCTTACTTGTCGTCATCGTCTTTGTAGTCAAATACATTGCTTAACGTTTTACC | 10.1252.X gDNA | pRab11-SAPIG0305-flag |
| FA40_BglII_307_F | CGAAAGATCTATGGAACTTGATGCATTAGTAATGC | 10.1252.X gDNA | pRab11-307-308-309 |
| FA101_EcoRI_309_R | CCAGAATTCTCAACTTCTTATATTATAATAATCTTG | 10.1252.X gDNA | pRab11-307-308-309 |
| FA218_pTrc_F | ATGGAACTTGATGCATTAGTAATG | pRab11-307-308-309 | pRab11-303-304-305flag-306-307-308-309 |
| FA219_pTrc_R | GGTCTGTTTCCTGTGTGAAATTG | pRab11-307-308-309 | pRab11-303-304-305flag-306-307-308-309 |
| FA220_303456_F | TTTCACACAGGAAACAGACCATGGGGGAAATAAAAGTTG | 10.1252.X gDNA | pRab11-303-304-305-306-307-308-309 |

| | | | |
|---|---|---|---|
| FA217_303456_rtrim | ACTAATGCATCAAGTTCCATTTATTATTCC | 10.1252.X gDNA | pRab11-303-304-305-306-307-308-309 |
| FA234_305flag_Q5F | GATGATGATAAATAAAAGGAGATTTAAAATGAATAATAC | pRab11-303-304-305-306-307-308-309 | pRab11-303-304-305flag-306-307-308-309 |
| FA235_305flag_Q5R | ATCTTTATAATCAAATACATTGCTTAACGTTTTAC | pRab11-303-304-305-306-307-308-309 | pRab11-303-304-305flag-306-307-308-309 |
| FA272_pRabflag_D306-9_Q5F | TATAAGAAGTTGAGAATTCTATC | pRab11-303-304-305flag-306-307-308-309 | pRab11-303-304-305flag |
| FA273_pRabflag_D306-9_Q5R | TTTAAATCTCCTTTTATTTATCATC | pRab11-303-304-305flag-306-307-308-309 | pRab11-303-304-305flag |
| FA301_D3078_F | TTATGGAATATTGAGGTGC | pRab11-303-304-305flag-306-307-308-309 | pRab11-303-304-305flag-306-309 |
| FA302_D3078_R | GTGTGTTTTGTAAATCTATATCATTTAG | pRab11-303-304-305flag-306-307-308-309 | pRab11-303-304-305flag-306-309 |
| FA293_p345flag_dsF | GAAGTTGAGAATTCTATCCATATG | pRab11-303-304-305flag | pRab11-303-304-305flag-307-308 |
| FA294_p345flag_dsR | CTCCTTTTATTTATCATCATCATC | pRab11-303-304-305flag | pRab11-303-304-305flag-307-308 |
| FA295_rbs3078_F | atgatgataaataaaaggagGTTTTTCACTGCTTTTATATTTTAAATTG | 10.1252.X gDNA | pRab11-303-304-305flag-307-308 |
| FA296_rbs3078_R | tggatagaattctcaacttcTCAATATTCCATAACTTTTACTTTAATATC | 10.1252.X gDNA | pRab11-303-304-305flag-307-308 |
| pRab_fwd | AATTCACTGGCCGTCGTTTTAC | pRab11 | pRab11-303-304-305flag-307 |
| pRab_rev | GTTAACGGTACCATCATACTCTATC | pRab11 | pRab11-303-304-305flag-307 |

| | | | |
|---|---|---|---|
| D308_fwd | agtatgatggtaccgttaacTATATTCAGGAGGTTTAGATCTATG | pRab11-303-304-305flag-307-308 | pRab11-303-304-305flag-307 |
| D308_rev | aaaacgacggccagtgaattTCATTCCAATTCATCCTC | pRab11-303-304-305flag-307-308 | pRab11-303-304-305flag-307 |
| pRab_fwd | AATTCACTGGCCGTCGTTTTAC | pRab11 | pRab11-303-304-305flag-308 |
| pRab_rev | GTTAACGGTACCATCATACTCTATC | pRab11 | pRab11-303-304-305flag-308 |
| D307part1_fwd | agtatgatggtaccgttaacTATATTCAGGAGGTTTAGATCTATG | pRab11-303-304-305flag-307-308 | pRab11-303-304-305flag-308 |
| D307part1_rev | aattgatttcTTATTTATCATCATCATCTTTATAATCAAATAC | pRab11-303-304-305flag-307-308 | pRab11-303-304-305flag-308 |
| D307part2_fwd | atgataaataaGAAATCAATTTAGATGAGGATG | pRab11-303-304-305flag-307-308 | pRab11-303-304-305flag-308 |
| D307part2_rev | aaaacgacggccagtgaattTCAATATTCCATAACTTTTACTTTAATATC | pRab11-303-304-305flag-307-308 | pRab11-303-304-305flag-308 |
| CM367 | CTCACTGAATTTCATTTTggcattagcgacagg | 10.1252.X gDNA | pRab11-Hlass-SAPIG0305CT |
| CM368 | cctgtcgctaatgccAAAATGAAATTCAGTGAG | 10.1252.X gDNA | pRab11-Hlass-SAPIG0305CT |
| CM369 | gcgcGAGCTCTTAAAATACATTGCTTAACGTTTT | 10.1252.X gDNA | pRab11-Hlass-SAPIG0305CT |
| CM286 | GCGC ggtacc AGG AGG TTT CTA GTT ATG aaa aca cgt ata gtc agc | 10.1252.X gDNA | pRab11-Hlass-SAPIG0305CT |
| FA54_NcoI_307_F | GCTGACCATGGAACTTGATGCATTAGTAATGCC | 10.1252.X gDNA | pTrc-307h |
| FA55_HindIII_His6307_R | TAAAGCTTTTAATGATGGTGATGATGGTGTTCCAATTCATCCTCATCTAAATTG | 10.1252.X gDNA | pTrc-307h |
| FA54_NcoI_307_F | GCTGACCATGGAACTTGATGCATTAGTAATGCC | 10.1252.X gDNA | pTrc-307-308h |
| FA57_HindIII_His6308_R | TAAAGCTTTTAATGATGGTGATGATGGTGATATTCCATAACTTTTACTTTAATATCG | 10.1252.X gDNA | pTrc-307-308h |

| | | | |
|---|---|---|---|
| FA108_Q5 308ts F | TTCAGGTGGTTCATCAGCTTGGTCACACCCACAATTCGAAAAATGAGGTACCCACGTGTCG | pTrc-307-308h | pTrc-307-308ts |
| FA109_Q5 308ts R | CCACCACCTGAACCACCACCTTTTTCGAATTGTGGGTGTGACCAATATTCCATAACTTTTACTTTAATATCGATCC | pTrc-307-308h | pTrc-307-308ts |
| FA220_303456_F | TTTCACACAGGAAACAGACCATGGGGGAAATAAAAGTTG | 10.1252.X gDNA | pTrc-303-304-305-306-307-308ts |
| FA216_303456_r | ACTAATGCATCAAGTTCCATTTATTATTCCTCCAATTTAAAATATAAAAG | 10.1252.X gDNA | pTrc-303-304-305-306-307-308ts |
| FA218_pTrc_F | ATGGAACTTGATGCATTAGTAATG | pTrc-307-308ts | pTrc-303-304-305-306-307-308ts |
| FA219_pTrc_R | GGTCTGTTTCCTGTGTGAAATTG | pTrc-307-308ts | pTrc-303-304-305-306-307-308ts |
| FA236_his305_Q5F | CACCACCACGGGAATAAAATAAAAATGTCAGAAG | pTrc-303-304-305-306-307-308ts | pTrc-303-304-h305-306-307ts |
| FA237_his305_Q5R | ATGATGATGCATTTTATACTCCTTTACTCTTTTATATTTATAATTG | pTrc-303-304-305-306-307-308ts | pTrc-303-304-h305-306-307ts |
| FA209_BamHI_305_F | GCCAGGATCCAGGGAATAAAATAAAAATGTCAGAAGTG | 10.1252.X gDNA | pACD-his305 |
| FA210_SalI_305_R | GCTTGTCGACTTAAAATACATTGCTTAACGTTTTACC | 10.1252.X gDNA | pACD-his305 |

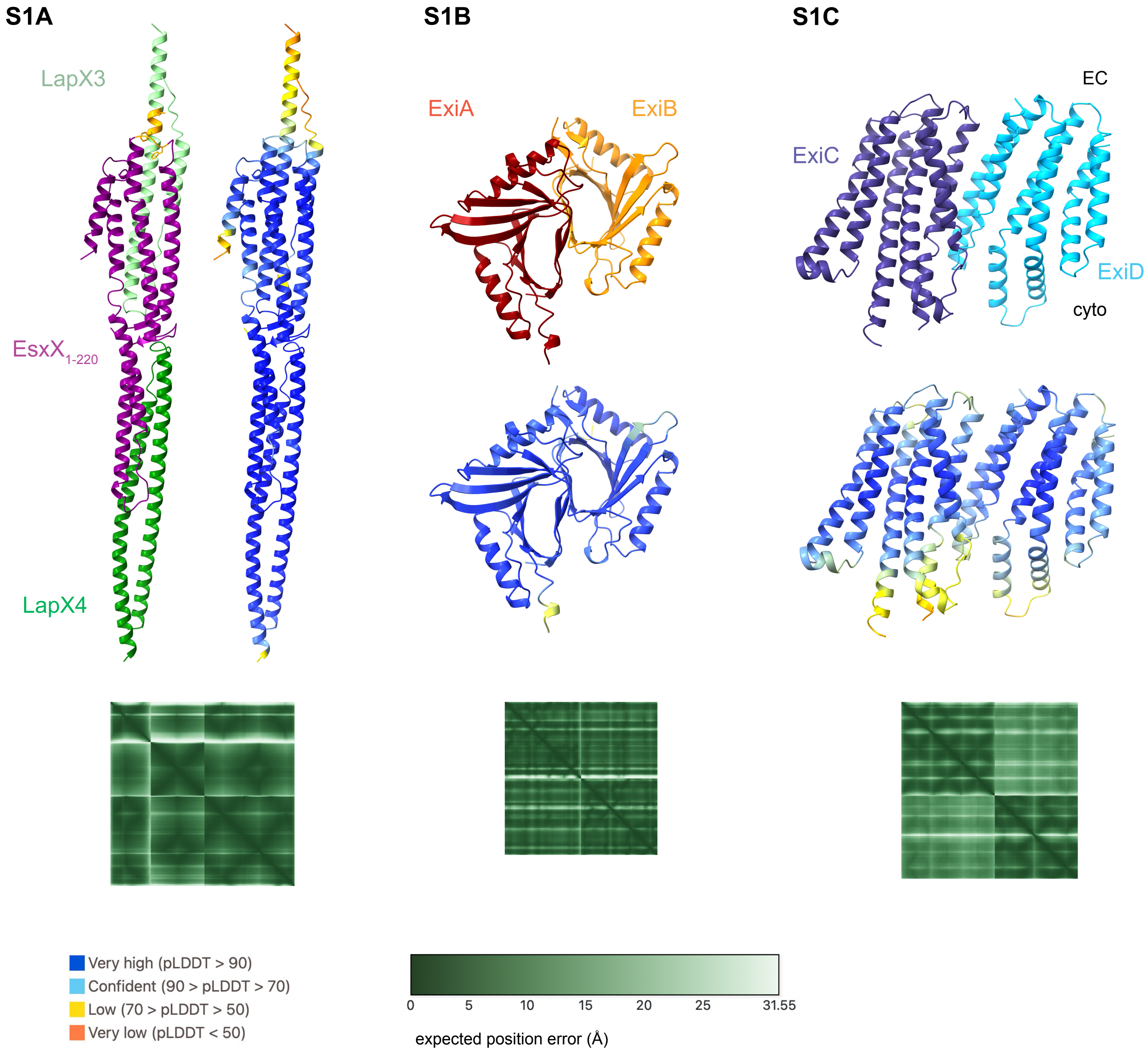
S1A
S1B
S1C
LapX3
EsxX1-220
LapX4
ExiA
ExiB
EC
ExiC
ExiD
cyto
Very high (pLDDT > 90)
Confident (90 > pLDDT > 70)
Low (70 > pLDDT > 50)
Very low (pLDDT < 50)
0
5
10
15
20
25
31.55
expected position error (Å)

**S2**

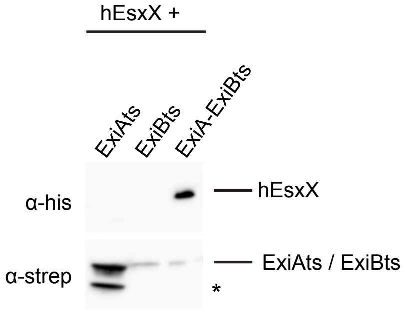
hEsxX +
ExiAts
ExiBts
ExiA-ExiBts
α-his
hEsxX
α-strep
ExiAts / ExiBts
*

S3A

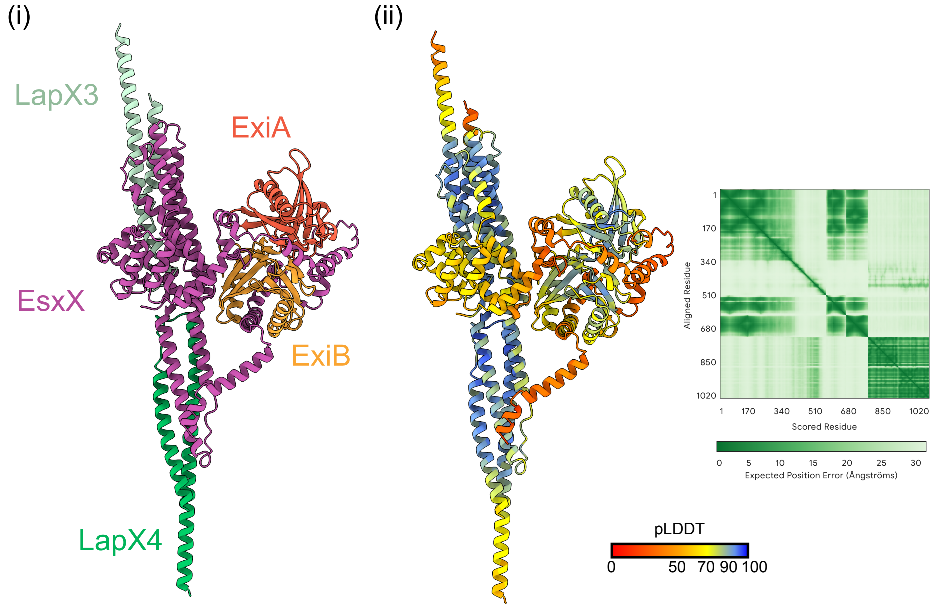
(i)
(ii)
LapX3
ExiA
EsxX
ExiB
LapX4
Aligned Residue
Scored Residue
Expected Position Error (Ångströms)
pLDDT
0
50 70 90 100

S3B

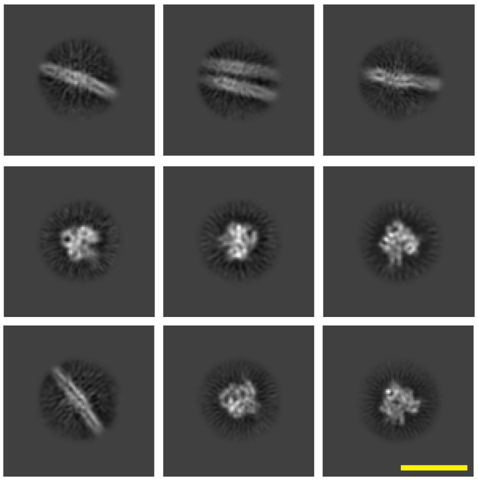

S3C

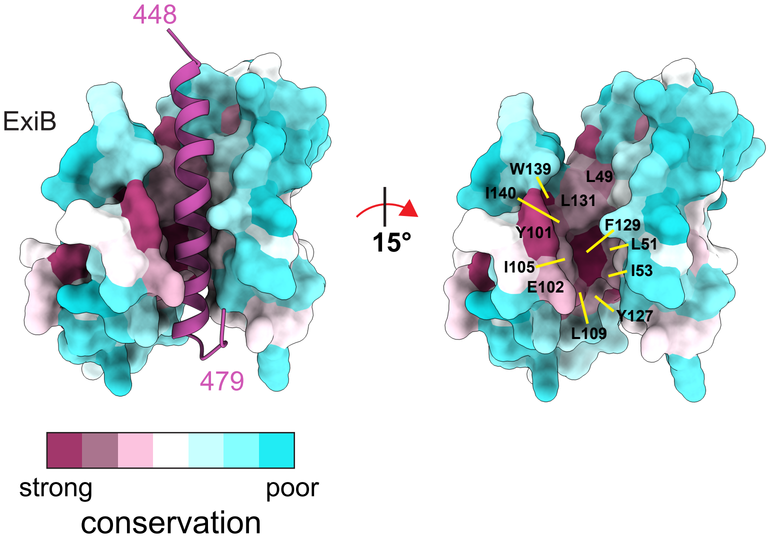
448
ExiB
479
15°
W139
L49
I140
L131
Y101
F129
L51
I105
I53
E102
Y127
L109
strong
poor
conservation

**S4**

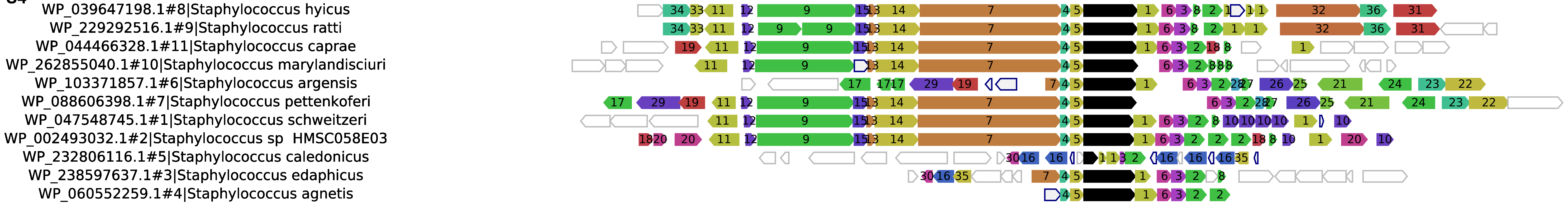
WP_039647198.1#8|Staphylococcus hyicus
WP_229292516.1#9|Staphylococcus ratti
WP_044466328.1#11|Staphylococcus caprae
WP_262855040.1#10|Staphylococcus marylandisciuri
WP_103371857.1#6|Staphylococcus argensis
WP_088606398.1#7|Staphylococcus pettenkoferi
WP_047548745.1#1|Staphylococcus schweitzeri
WP_002493032.1#2|Staphylococcus sp HMSC058E03
WP_232806116.1#5|Staphylococcus caledonicus
WP_238597637.1#3|Staphylococcus edaphicus
WP_060552259.1#4|Staphylococcus agnetis

**S5**

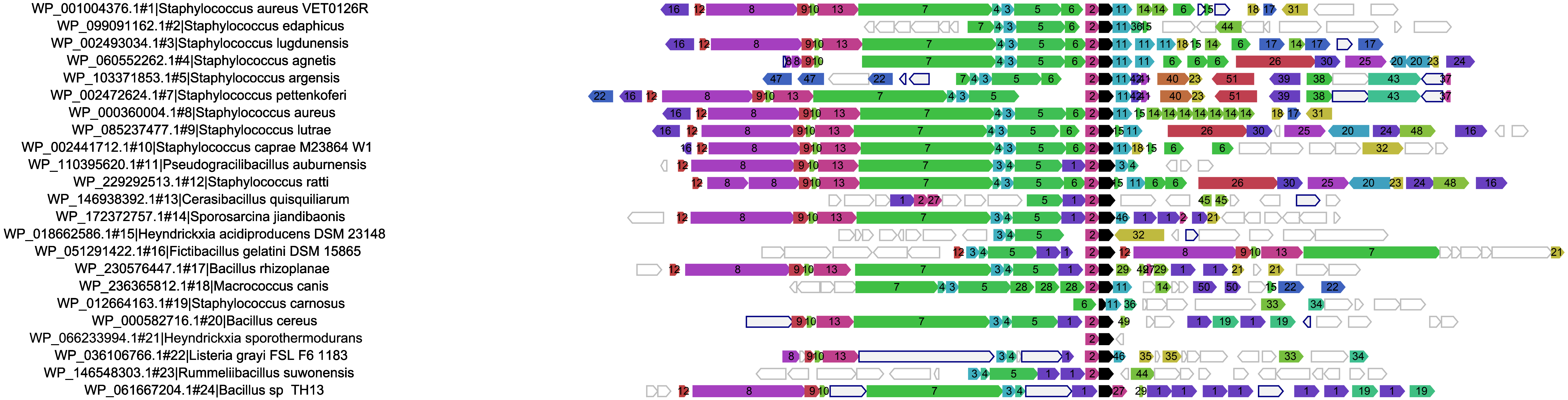
WP_001004376.1#1|Staphylococcus aureus VET0126R
WP_099091162.1#2|Staphylococcus edaphicus
WP_002493034.1#3|Staphylococcus lugdunensis
WP_060552262.1#4|Staphylococcus agnetis
WP_103371853.1#5|Staphylococcus argensis
WP_002472624.1#7|Staphylococcus pettenkoferi
WP_000360004.1#8|Staphylococcus aureus
WP_085237477.1#9|Staphylococcus lutrae
WP_002441712.1#10|Staphylococcus caprae M23864 W1
WP_110395620.1#11|Pseudogracilibacillus auburnensis
WP_229292513.1#12|Staphylococcus ratti
WP_146938392.1#13|Cerasibacillus quisquiliarum
WP_172372757.1#14|Sporosarcina jiandibaonis
WP_018662586.1#15|Heyndrickxia acidiproducens DSM 23148
WP_051291422.1#16|Fictibacillus gelatini DSM 15865
WP_230576447.1#17|Bacillus rhizoplanae
WP_236365812.1#18|Macrococcus canis
WP_012664163.1#19|Staphylococcus carnosus
WP_000582716.1#20|Bacillus cereus
WP_066233994.1#21|Heyndrickxia sporothermodurans
WP_036106766.1#22|Listeria grayi FSL F6 1183
WP_146548303.1#23|Rummeliibacillus suwonensis
WP_061667204.1#24|Bacillus sp TH13

**S6**

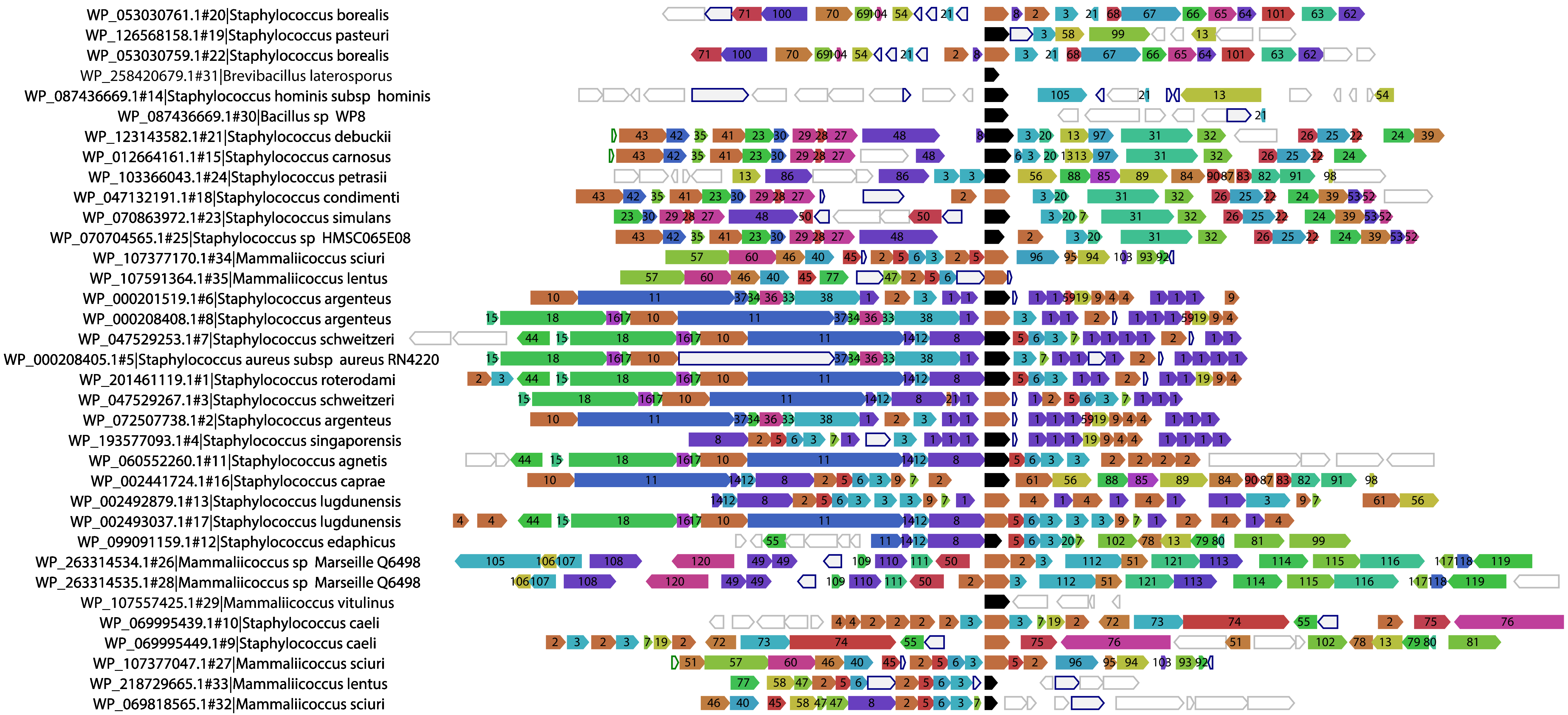
WP_053030761.1#20|Staphylococcus borealis
WP_126568158.1#19|Staphylococcus pasteuri
WP_053030759.1#22|Staphylococcus borealis
WP_258420679.1#31|Brevibacillus laterosporus
WP_087436669.1#14|Staphylococcus hominis subsp hominis
WP_087436669.1#30|Bacillus sp WP8
WP_123143582.1#21|Staphylococcus debuckii
WP_012664161.1#15|Staphylococcus carnosus
WP_103366043.1#24|Staphylococcus petrasii
WP_047132191.1#18|Staphylococcus condimenti
WP_070863972.1#23|Staphylococcus simulans
WP_070704565.1#25|Staphylococcus sp HMSC065E08
WP_107377170.1#34|Mammaliicoccus sciuri
WP_107591364.1#35|Mammaliicoccus lentus
WP_000201519.1#6|Staphylococcus argenteus
WP_000208408.1#8|Staphylococcus argenteus
WP_047529253.1#7|Staphylococcus schweitzeri
WP_000208405.1#5|Staphylococcus aureus subsp aureus RN4220
WP_201461119.1#1|Staphylococcus roterodami
WP_047529267.1#3|Staphylococcus schweitzeri
WP_072507738.1#2|Staphylococcus argenteus
WP_193577093.1#4|Staphylococcus singaporensis
WP_060552260.1#11|Staphylococcus agnetis
WP_002441724.1#16|Staphylococcus caprae
WP_002492879.1#13|Staphylococcus lugdunensis
WP_002493037.1#17|Staphylococcus lugdunensis
WP_099091159.1#12|Staphylococcus edaphicus
WP_263314534.1#26|Mammaliicoccus sp Marseille Q6498
WP_263314535.1#28|Mammaliicoccus sp Marseille Q6498
WP_107557425.1#29|Mammaliicoccus vitulinus
WP_069995439.1#10|Staphylococcus caeli
WP_069995449.1#9|Staphylococcus caeli
WP_107377047.1#27|Mammaliicoccus sciuri
WP_218729665.1#33|Mammaliicoccus lentus
WP_069818565.1#32|Mammaliicoccus sciuri

**S7**

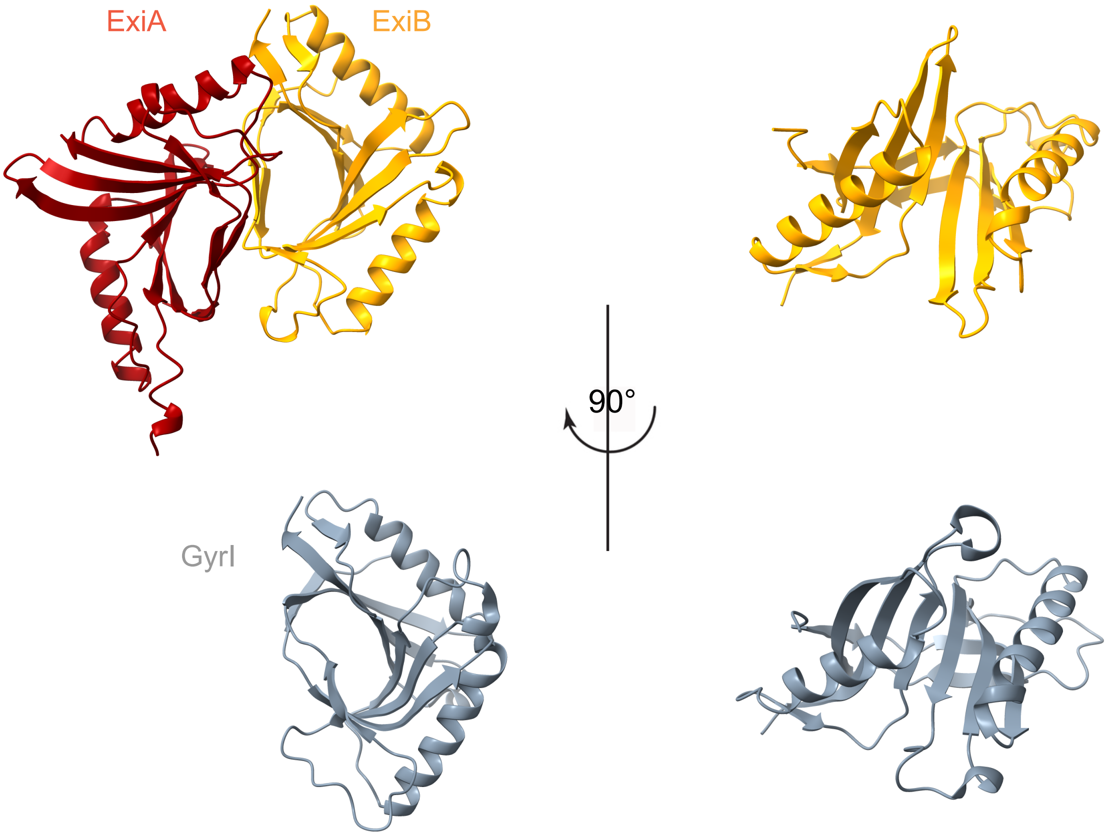
